# Gut bacterial diversity and physiological traits of *Anastrepha fraterculus* Brazilian-1 morphotype males are affected by antibiotic treatment

**DOI:** 10.1101/360693

**Authors:** M. Laura Juárez, Lida E. Pimper, Guillermo E. Bachmann, Claudia A. Conte, M. Josefina Ruiz, Lucía Goane, Pilar Medina Pereyra, Felipe Castro, Julieta Salgueiro, Jorge L. Cladera, Patricia E. Fernández, Kostas Bourtzis, Silvia B. Lanzavecchia, M. Teresa Vera, Diego F. Segura

**Affiliations:** Cátedra Terapéutica Vegetal, Facultad de Agronomía y Zootecnia (FAZ), Universidad Nacional de Tucumán (UNT), Tucumán, Argentina; Unidad Ejecutora Lillo, Fundación Miguel Lillo, Tucumán, Argentina; Consejo Nacional de Investigaciones Científicas y Técnicas (CONICET); Instituto de Genética Ewald A. Favret (IGEAF), Instituto Nacional de Tecnología Agropecuaria (INTA), Castelar, Argentina; Instituto de Fisiología Animal, Fundación Miguel Lillo, Tucumán, Argentina; Estaciòn Agropecuaria Delta, Instituto Nacional de Tecnología Agropecuaria (INTA), Argentina; Insect Pest Control Laboratory, Joint FAO/IAEA Division of Nuclear Techniques in Food and Agriculture, Vienna, Austria

**Keywords:** South American fruit fly, symbiont, antibiotics, nutritional reserves, survival, Sterile Insect Technique

## Abstract

**Background:** The interaction between gut bacterial symbionts and Tephritidae became the focus of several studies that showed that bacteria contributed to the nutritional status and the reproductive potential of its fruit fly hosts. *Anastrepha fraterculus* is an economically important fruit pest in South America. This pest is currently controlled by insecticides, which prompt the development of environmentally friendly methods such as the sterile insect technique (SIT). For SIT to be effective, a deep understanding of the biology and sexual behavior of the target species is needed. Although many studies have contributed in this direction, little is known about the composition and role of *A. fraterculus* symbiotic bacteria. In this study we tested the hypothesis that gut bacteria contribute to nutritional status and reproductive success of *A. fraterculus* males.

**Methods:** Wild and laboratory-reared males were treated with antibiotics (AB) and provided sugar (S) or sugar plus protein (S+P) as food sources. The effect of AB on the gut bacteria diversity was assessed through DGGE and sequencing of the V6-V9 variable region of the bacterial 16S *rRNA* gene.

**Results:** AB affected the bacterial community of the digestive tract of *A. fraterculus*, in particular bacteria belonging to the Enterobacteriaceae family, which was the dominant bacterial group in the control flies (i.e., non-treated with AB). AB negatively affected parameters directly related to the mating success of laboratory males and their nutritional status. AB also affected males’ survival under starvation conditions. The effect of AB on the behaviour and nutritional status of the males depended on two additional factors: the origin of the males and the presence of a proteinaceous source in the diet.

**Conclusions:** Our results suggest that *A. fraterculus* males gut contain symbiotic organisms that are able to exert a positive contribution on *A. fraterculus* males’ fitness, although the physiological mechanisms still need further studies.

## Background

Insects maintain a close and complex association with microbial communities, ranging from parasitic relationships to commensalism and obligate mutualism [1, 2]. The contributions of gut bacteria to their insect hosts are diverse [see 3 for a review], but probably the most important is associated to its nutrition. Insects use the metabolic pathways of bacteria to obtain nutritional resources otherwise unavailable and thus are able to survive on suboptimal or nutrient-poor diets [3–6]. Bacterial symbionts have also been shown to have a protective function of their insect hosts, to the point that are considered to act as an additional immune system [4, 7, 8]. Although the way that this occurs is still unknown in most cases [3], Brownlie and Johnson [8] describe the production of toxins or antibiotics by gut bacteria that would protect the host against pathogens. Other benefits include improving digestion efficiency, the acquisition of digestive enzymes, some of them associated with detoxification, and the provision of vitamins, nitrogen, specific amino acids and carbon [4]. Bacterial symbionts have also been shown to contribute with chemical compounds that participate in the communication between the hosts and other individuals, present either in the volatiles emitted or retained in the insect cuticle [3, 4, 9]. Moreover, the presence of gut bacteria has been associated to the improvement of developmental and reproductive parameters, such as mating behavior [3, 10].

The study of the interactions that bacteria and their hosts establish has followed different experimental approaches [6]. One of these approaches is to phenotypically characterize the bacterial community present in the gut by culture-dependent techniques or to determine its function inferred from their genome sequence by culture-independent molecular methods [11–18]. Another indirect way to assess the effect of gut bacteria is to evaluate the effect of adding antibiotics (AB) into the insect diets and compare parameters associated to the fitness of AB-treated and non-treated insects [5, 19–23]. Alternatively, other studies have taken a more direct approach in which insects were fed specific bacterial species to determine potential benefits associated with the increase of bacteria titers in their gut [10, 24–31].

The sterile insect technique (SIT) is an environmentally friendly and species-specific control method commonly used against tephritid fruit fly pests. The SIT consists of mass production, sterilization, and release of males to mate with wild females [32, 33]. For an effective implementation of the SIT, a deep understanding of the biology of the targeted species is needed, particularly its sexual behavior [33]. Thus, the interaction between gut bacteria and fruit flies has become the focus of several studies in recent years. Combining traditional microbiological methods and molecular techniques, the composition of the bacterial community associated to Tephritidae fruit flies has been characterized for some species. Studies on *Ceratitis capitata* Wiedemann, the Mediterranean fruit fly, showed that gut bacterial community is comprised mainly by members of the family Enterobacteriaceae [10, 12, 34, 35]. However, the monophagous olive fruit fly *Bactrocera oleae* Gmelin is characterized by the presence of the obligatory symbiont *Candidatus* Erwinia dacicola that colonize a specialized evagination of the digestive tract while in the gut a limited number of the bacterial species have been reported such as *Acetobacter tropicalis* [36–38]. Through indirect (AB treatment) or direct (feeding larvae or adults) approaches, gut bacteria were shown to contribute to several biological parameters of their hosts, such as longevity [20, 22, 27], fecundity [5, 21, 29], development, productivity and mating success [10, 19, 25, 27, 30, 31, 39]. The South American fruit fly, *Anastrepha fraterculus* Wiedemann (Diptera: Tephritidae), is a major pest causing considerable damage to a wide spectrum of host fruit species, many of them of economic importance [40, 41]. Currently, the only control method for this species is through the use of insecticides which prompt the development of alternative control methods such as the SIT. The efficacy of the technique depends on the mating success of males released in the field. Many studies so far have provided valuable information in this regard [42–49]. However, despite the important role that gut bacteria have on the development, productivity and the reproductive success of other Tephritidae flies, no study addressed the significance of these interactions for *A. fraterculus* so far. Because understanding how bacterial symbionts affect the overall fitness of sterile males may contribute to the efficacy of the SIT, in the present study, and as an initial approach, we tested the hypothesis that gut bacteria contribute to nutritional and reproductive aspects of wild and laboratory-reared *A. fraterculus* males from the Brazilian-1 morphotype. Following an indirect approach, we tested the effect of AB treatment on several parameters associated to males’ reproductive success such as male sexual performance, and sexual communication mediated by chemical signals and behavioral displays. Also, the nutritional status and the starvation resistance of AB treated and non-treated males were evaluated. In parallel, the effect of AB on the gut bacteria diversity was assessed through molecular techniques. As previous studies in other species have shown that the dietary regime, particularly the protein content of the adult diet, interacts with the presence of gut bacteria, we carried out the above experiments providing a complete diet (sugar and a protein source) and a nutritionally poor diet that contained only sugar.

## Results

### Diet consumption

The presence of AB had no impact on diet consumption, irrespectively of the origin of the flies or the diet given (F_1,2_ = 0.02, P = 0.9107 for S fed lab males; F_1,2_ = 6.52, P = 0.1252 for S+P fed lab males; F_1,2_ = 1.35, P = 0.3655 for S fed wild males; F_1,2_ =0.10, P = 0.7776 for S+P fed wild males) (Fig. 1).

**Figure 1.**
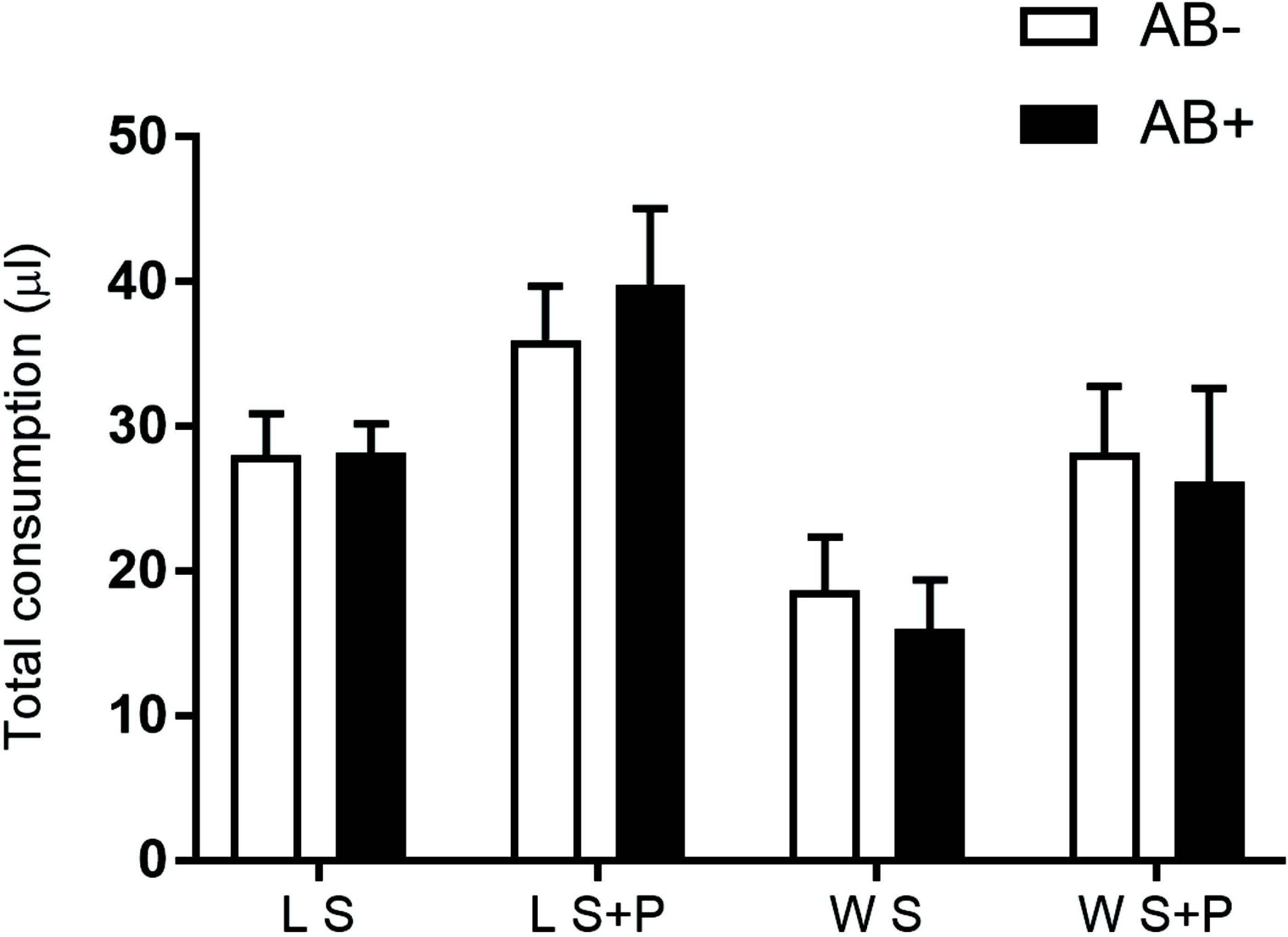
Effect of antibiotics treatment on laboratory and wild *Anastrepha fraterculus* males’ consumption. Individual total consumption (µl) of males exposed to two different diets with or without the antibiotic addition (AB): S and S+AB diets, or S+P and S+P+AB in a dual choice experiment.

### Molecular characterization of gut bacteria

Total DNA from single *A. fraterculus* guts was used to describe the bacterial community associated to male flies from different origin, types of food and AB treatment using molecular tools. The V6-V9 region of the bacterial 16S *rRNA* gene was amplified by PCR using universal primers. 27 bands of approximately 420 bp were excised from the DGGE gels, and 14 PCR fragments were successfully sequenced to identify the associated bacterial taxonomic groups. The nucleotide sequences obtained for the rest of the PCR products (13) presented double peaks and low quality, showing the potential presence of several amplicons in the same sample. The results of differential band sequencing obtained from the different combinations of treatments showed the presence of microorganisms closely related to the Proteobacteria, distributed as: Gamaproteobacteria, 71% and Alphaproteobacteria, 29% of the total bands (Table 1, Additional files 1; Fig. S1). The use of both distance matrix (Fig. 2) and character-based (parsimony, data not shown) methods resulted in the construction of similar phylogenetic trees. All bacterial strains were phylogenetically related to taxonomic groups of Proteobacteria (linked to Enterobactereales, Xanthomonadales and Alphaproteobacteria class) (Fig. 2), in accordance with the closest relatives found using RDP/Blast search (Table 1). The analysis of the sequences revealed that the Enterobacteriaceae family is the dominant bacterial group in the *A. fraterculus* gut, in both wild and lab flies (S or S+P diet). AB treated flies (wild and lab) fed with a S+P diet contained species of the genus *Stenotrophomonas* sp., and Alphaproteobacteria class; whereas AB treated flies (wild and lab) fed with sugar contained only species of the Alphaproteobacteria class (Table 1; Fig. 2).

**Figure 2.**
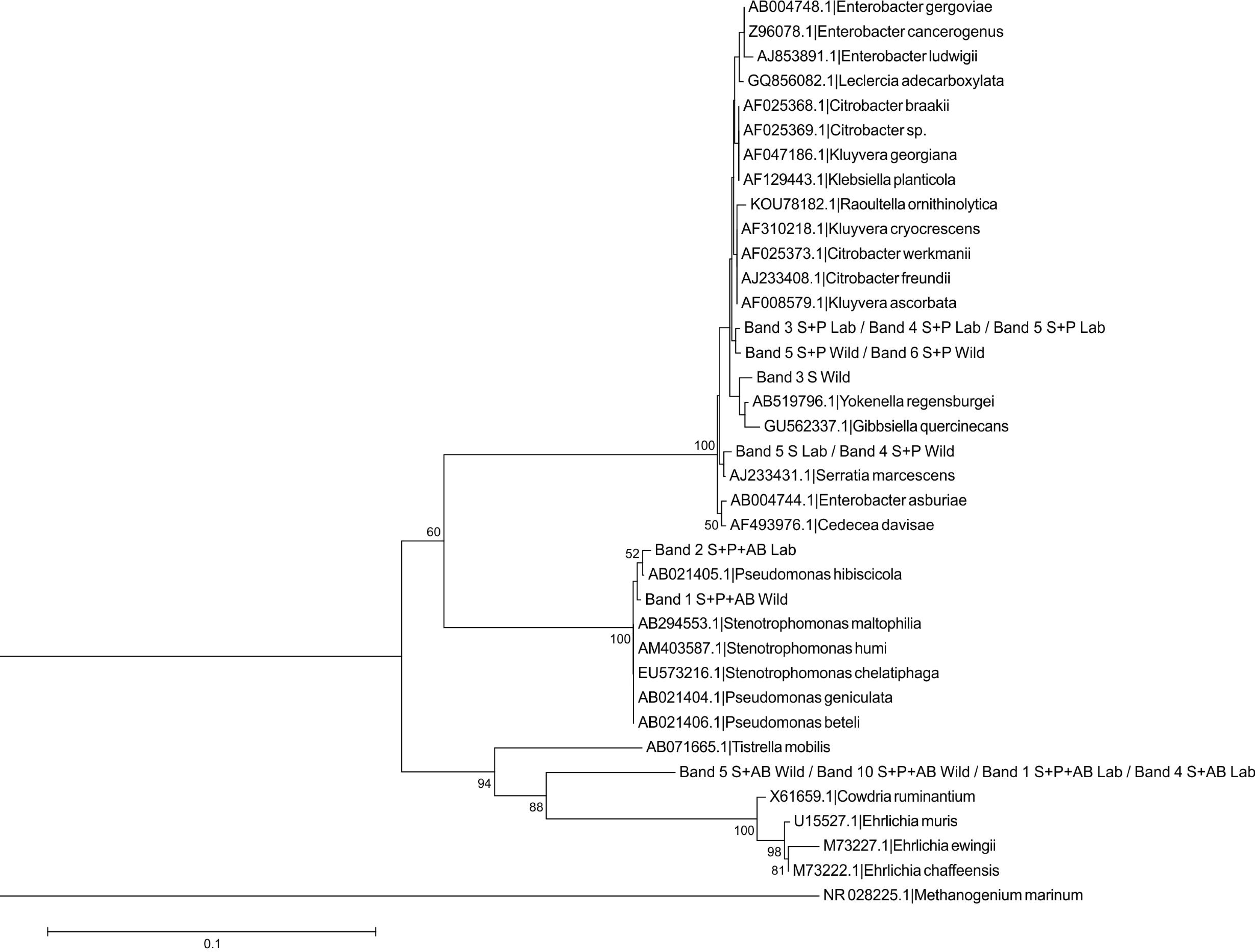
Phylogenetic tree based on V6-V9 16S *rRNA* gene sequence analysis of *A. fraterculus* gut bacteria and the closest relative taxa. The tree is based on Neighbor-Joining method (Jukes-Cantor distance), using a 50% conservation filter. Numbers on the nodes present % bootstrap values based on 1000 replicates. Scale bar indicates 10% estimated sequence divergence. The 16S *rRNA* gene sequences of *Methanogenium marinum* were arbitrarily chosen as an outgroup.

**Table 1.** Analysis of V6-V9 16S *rRNA* gene sequences obtained from DGGE profiles and sequencing.

| DGGE Band Number | Taxonomic group (RDP) | Closest related sequence (BLAST)<br>(Genbank Accession number) | Nucleotide<br>bases<br>compared | Similarity |
| --- | --- | --- | --- | --- |
| 3 S+P Lab / 4 S+P Lab /<br>5 S+P Lab | Gamaproteobacteria/ Enterobacteriales/<br>Enterobacteriaceae/<br>unclassified_Enterobacteriaceae | *Raoultella planticola* strain Ns8<br>(MG544105.1)<br>*Serratia marcescens* subsp.<br>*marcescens* (HG326223.1 ) | 389 | 99% |
| 4 S+P Wild / 5 S Lab | Gamaproteobacteria/ Enterobacteriales/<br>Enterobacteriaceae/<br>unclassified_Enterobacteriaceae | *Enterobacter soli* strain YHBG2<br>(MG516168.1)<br>*Serratia marcescens* subsp.<br>*marcescens* (MG516113.1) | 384 | 100% |
| 1 S+P+AB Wild /<br>2 S+P+AB Lab | Gamaproteobacteria/ Xanthomonadales/<br>Xanthomonadaceae/ *Stenotrophomonas* | *Stenotrophomonas maltophilia*<br>(MG546679.1)<br>*Stenotrophomonas maltophilia*<br>(MG546678.1) | 388 | 100% |
| 5 S+P Wild / 6 S+P Wild | Gamaproteobacteria/ Enterobacteriales/<br>Enterobacteriaceae/ unclassified<br>_Enterobacteriaceae | Uncultured *Enterobacter* sp. clone 03<br>(KJ526996.1)<br>*Klebsiella aerogenes* strain JMB006<br>(MG546216.1) | 387 | 99% |
| 10 S+P+AB Wild /<br>5 S+AB Wild / 1 S+P+AB<br>Lab / 4 S+AB Lab | Alphaproteobacteria/ unclassified_Alpha-<br>proteobacteria | Uncultured alpha proteobacterium<br>(HM111616.1) | 390 | 99% |
| 3 S Wild | Gamaproteobacteria/Enterobacteriales/<br>Enterobacteriaceae/Citrobacter | *Klebsiella* sp. M5a1 (CP020657.1)<br>*Klebsiella oxytoca* strain FCX2 16S<br>(KU942497.1) | 387 | 100% |

### Male mating competitiveness

Overall, the mean percentage of copulations achieved by wild males was not affected by AB (F_1,2_ = 0.16, P = 0.7299 for S fed males; F_1,4_ = 1.31, P = 0.3163 for S+P fed males). In contrast, for lab males, the effect of AB depended on the diet. AB had a significantly negative impact on percentage of mating for S+P fed males (F_1,3_ = 18.71, P = 0.0228) while for males fed with S diet, the differences were not significant (F_1,2_ = 0.46, P = 0.5689) (Fig. 3A). Latency to mate was not significantly affected by AB neither for wild (W = 366.5, P = 0.1590 for S fed males; W = 4814.5, P = 0.1000 for S+P fed males) nor for lab males (W = 2762, P = 0.5256 for S fed males; W = 3857.5 P = 0.9155 for S+P fed males) (Fig. 3B). Copula duration was also not significantly affected by AB (F1,107 = 1.29, P = 0.2587 for S fed lab males; F1,128 = 0.12, P = 0.7291 for S+P fed lab males; F_1,36_ = 1.67, P = 0.2048 for S fed wild males; F1,128 = 0.90, P = 0.3441 for S+P fed wild males) (Fig. 3C).

**Figure 3.**
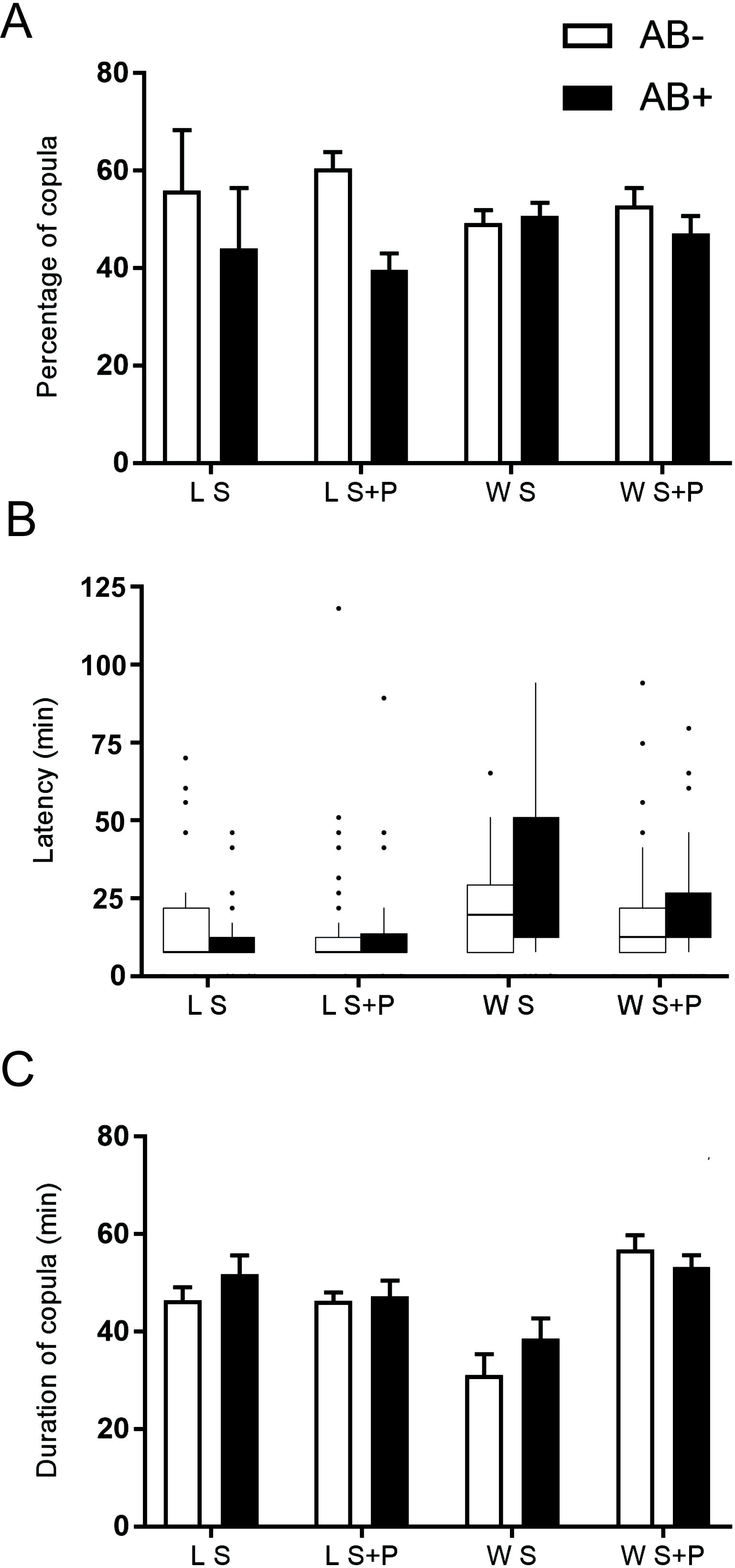
Effect of antibiotics treatment on laboratory and wild *Anastrepha fraterculus* male mating competitiveness. (A) Percentage of matings (B) Latency to copulate (time elapsed before copulation started) and (C) Duration of copula obtained by males fed with two different diets with or without addition of antibiotic (AB).

### Male calling behavior

Behavioral recordings showed that for S fed males, AB affected the mean number of wing fanning and salivary gland exposure (t = 2.148, d.f. = 14, p = 0.024; and t = 1.870, d.f. = 14, p = 0.041, respectively). For the two variables, males without AB performed these courtship-related behaviors more frequently than AB males (Fig. 4A, B). On the other hand, AB did not affect wing fanning or gland exposure in S+P fed males (t = 0.100, d.f. = 14, p = 0.461; and t = 0.387, d.f. = 14, p = 0.352, respectively) (Fig. 4A, B).

**Figure 4.**
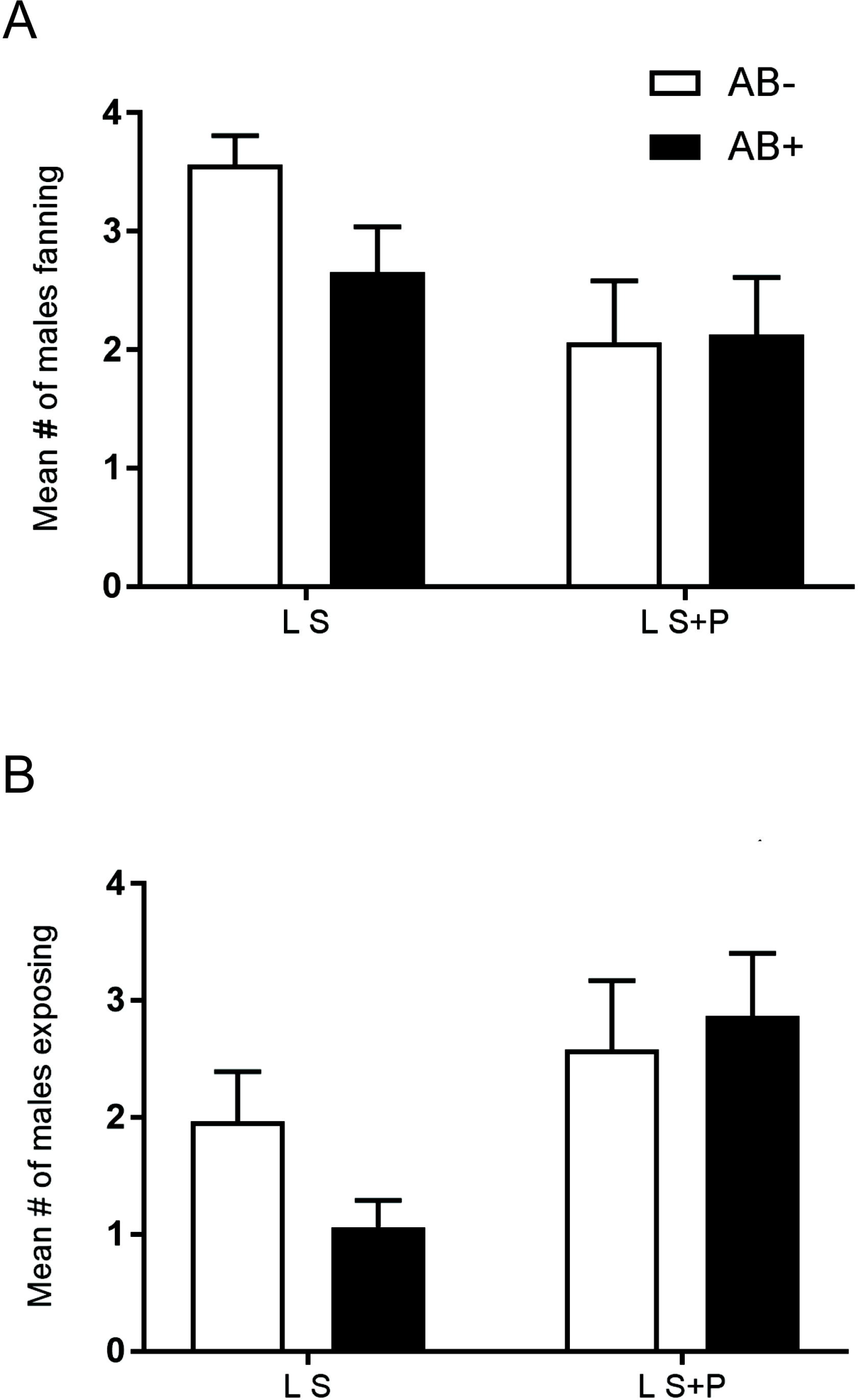
Effect of antibiotics treatment on laboratory *Anastrepha fraterculus* male calling behavior and pheromone release. (A) Number of males fed on S or S+AB and S+P or S+P+AB diets that were detected fanning their wings across the observational period. (B) Number of males fed on S or S+AB and S+P or S+P+AB diets that were detected exposing their salivary glands across the observational period.

### Male volatile and cuticle compounds

Ten compounds were quantified in the volatile collections of *A. fraterculus* males. For S fed males, we detected significantly higher amounts of three compounds (E-E-α-farnesene, anastrephin, epianastrephin) in the volatiles’ collections compared to S+AB fed males, whereas the remaining seven compounds showed no significant differences (Table 2). For males S+P males, no significant differences were detected for any of the 10 compounds between AB treated and non-treated males (Table 2). When antennally active compounds were combined, S fed males that were treated with AB released significantly less amount of these compounds than non-treated males whereas no differences between treated and non-treated males were detected for S+P males (Table 2).

**Table 2.** Relative abundances (mean ± S.E.) of compounds detected in the volatile collection of *Anastrepha fraterculus* males fed on S or S+P diets (N = 8). Results are shown as mean ± SE for AB treated and non-treated males and compared by means of a Student’s *t*-test.

| Ret. time (min) | Compound | KI | KI lit.<sup>c</sup> | Sugar fed males |  |  | Sugar + protein fed males |  |  |
| --- | --- | --- | --- | --- | --- | --- | --- | --- | --- |
|  |  |  |  | S | S+AB | p | S+P | S+P+AB | p |
| 10.76 | limonene | 1027 | 1031 | 0.06 ± 0.004 | 0.07 ± 0.004 | 0.45 | 0.19 ± 0.05 | 0.15 ± 0.005 | 0.27 |
| 10.94 | indane | 1033 | 1034 | 0 | 0 | - | 1.622 ± 0.36 | 1.34 ± 0.026 | 0.27 |
| 11.00 | unknown | 1035 | - | 0.01 ± 0.002 | 0.05 ± 0.031 | 0.18 | 0 | 0 | - |
| 11.40 | E-β-ocimene<sup>b</sup> | 1049 | 1050 | 1.84 ± 0.247 | 1.68 ± 0.215 | 0.32 | 6.554 ± 0.16 | 6.15 ± 0.098 | 0.41 |
| 14.00 | 4-Methylindane | 1141 | 1142 | 0 | 0 | - | 3.375 ± 0.08 | 2.89 ± 0.052 | 0.30 |
| 19.90 | suspensolide<sup>a</sup> | 1496 | no data | 0.42 ± 0.057 | 0.33 ± 0.058 | 0.16 | 0.543 ± 0.01 | 0.79 ± 0.018 | 0.15 |
| 19.92 | Z-E-α-farnesene<sup>b</sup> | 1498 | 1497 | 0.56 ± 0.012 | 0.39 ± 0.052 | 0.08 | 0.997 ± 0.03 | 1.11 ± 0.033 | 0.39 |
| 20.06 | E-E-α-farnesene<sup>ab</sup> | 1510 | 1508 | 9.21 ± 1.497 | 6.04 ± 0.818 | 0.04 | 14.996 ± 0.38 | 15.71 ± 0.447 | 0.45 |
| 20.98 | anastrephin<sup>a</sup> | 1596 | no data | 0.31 ± 0.037 | 0.21 ± 0.038 | 0.04 | 0.571 ± 0.01 | 0.78 ± 0.025 | 0.24 |
| 21.12 | epianastrephin<sup>ab</sup> | 1610 | 1621 | 0.91 ± 0.128 | 0.59 ± 0.071 | 0.04 | 1.498 ± 0.03 | 2.21 ± 0.072 | 0.19 |
| | Sum of EAG+ compounds | - | - | 11.93 ± 1.712 | 8.28 ± 1.09 | 0.05 | 23.106 ± 5.43 | 23.73 ± 5.641 | 0.23 |

a Compound identified by comparison with authentic standards.
b Compound that triggers a positive EAG response in female's antennae (Brizova et al. 2013; Bachmann 2016).
c: KI: Kovats index obtained for a DB5 / HP5 column and similar chromatographic conditions from www.pherobase.com and webbook.nist.gov.

Fifteen compounds were quantified in the cuticle extracts of *A. fraterculus* males. We did not detect significant differences between AB treated and non-treated males in any compound for any of the two diets (Table 3). The same result was found when antennally active compounds were added (Table 3).

**Table 3.** Relative abundances (mean ± S.E.) of compounds detected in the cuticle extracts of *Anastrepha fraterculus* males fed on S or S+P diets (N = 8). Results are shown as mean ± SE for AB treated and non-treated males and compared by means of a Student’s *t-*test.

| Ret. time (min) | Compound | KI | KI lit.<sup>c</sup> | Sugar fed males |  |  | Sugar + protein fed males |  |  |
| --- | --- | --- | --- | --- | --- | --- | --- | --- | --- |
|  |  |  |  | S | S+AB | p | S+P | S+P+AB | p |
| 19.90 | suspensolide<sup>a</sup> | 1496 | no data | 0.11 ± 0.001 | 0.12 ± 0.002 | 0.31 | 0.07 ± 0.002 | 0.06 ± 0.001 | 0.33 |
| 20.06 | E-E-α-farnesene<sup>ab</sup> | 1510 | 1508 | 0.60 ± 0.014 | 0.44 ± 0.007 | 0.16 | 0.57 ± 0.035 | 0.91 ± 0.053 | 0.31 |
| 20.98 | anastrephin<sup>a</sup> | 1596 | no data | 0.11 ± 0.002 | 0.12 ± 0.004 | 0.43 | 0.10 ± 0.006 | 0.27 ± 0.019 | 0.22 |
| 21.12 | epianastrephin<sup>ab</sup> | 1610 | 1621 | 0.55 ± 0.013 | 0.52 ± 0.016 | 0.44 | 0.46 ± 0.026 | 1.00 ± 0.066 | 0.25 |
| 23.79 | nonadecane* | 1900 | 1900 | 0.47 ± 0.011 | 0.45 ± 0.006 | 0.43 | 0.33 ± 0.008 | 0.66 ± 0.032 | 0.18 |
| 24.46 | monounsaturated alkene (C<sub>20</sub>) | 1981 | - | 0.11 ± 0.002 | 0.11 ± 0.003 | 0.47 | 0.12 ± 0.004 | 0.29 ± 0.016 | 0.18 |
| 25.23 | monounsaturated alkene (C<sub>21</sub>) | 2079 | - | 1.02 ± 0.029 | 1.12 ± 0.028 | 0.40 | 0.78 ± 0.022 | 1.87 ± 0.094 | 0.16 |
| 25.28 | monounsaturated alkene (C<sub>21</sub>) | 2084 | - | 14.24 ± 0.417 | 15.16 ± 0.376 | 0.44 | 19.24 ± 0.621 | 44.65 ± 2.460 | 0.18 |
| 25.44 | heneicosane<sup>a</sup> | 2100 | 2100 | 20.53 ± 0.431 | 20.94 ± 0.352 | 0.48 | 10.50 ± 0.344 | 9.87 ± 0.245 | 0.45 |
| 26.23 | monounsaturated alkene (C<sub>22</sub>) | 2180 | - | 0.81 ± 0.025 | 0.86 ± 0.011 | 0.44 | 0.64 ± 0.022 | 1.10 ± 0.044 | 0.20 |
| 27.32 | monounsaturated alkene (C<sub>23</sub>) | 2274 | - | 1.96 ± 0.061 | 2.11 ± 0.037 | 0.42 | 0.64 ± 0.016 | 1.33 ± 0.051 | 0.13 |
| 27.40 | monounsaturated alkene (C<sub>23</sub>) | 2280 | - | 22.89 ± 0.066 | 21.83 ± 0.346 | 0.44 | 18.59 ± 0.473 | 34.27 ± 1.600 | 0.20 |
| 27.64 | tricosane<sup>a</sup> | 2300 | 2300 | 3.40 ± 0.582 | 3.57 ± 0.058 | 0.44 | 1.65 ± 0.047 | 1.81 ± 0.053 | 0.42 |
| 29.20 | tetracosane<sup>a</sup> | 2400 | 2400 | 2.49 ± 0.029 | 2.25 ± 0.022 | 0.27 | 1.04 ± 0.023 | 0.74 ± 0.016 | 0.18 |
| 31.25 | pentacosane<sup>a</sup> | 2500 | 2500 | 3.28 ± 0.058 | 3.23 ± 0.040 | 0.48 | 1.62 ± 0.038 | 1.64 ± 0.039 | 0.48 |
| | Sum of EAG + compounds | - | - | 1.15 ± 0.021 | 0.96 ± 0.020 | 0.50 | 1.03 ± 0.032 | 1.91 ± 0.024 | 0.38 |

a Compound identified by comparison with authentic standards.
b Compound that triggers a positive EAG response in female's antennae (Brizova et al. 2013; Bachmann 2016).
c: KI: Kovats index obtained for a DB5 / HP5 column and similar chromatographic conditions from www.pherobase.com and webbook.nist.gov.

### Starvation resistance

Laboratory males fed on S and treated with AB lived longer under starvation than S-fed non-treated males (χ^2^ = 5.28, p = 0.0215). For S+P males, AB treatment had no effect (χ^2^ = 2.28, p = 0.1311) (Fig. 5A). Conversely, S fed wild males treated with AB lived less than non-treated males (χ^2^ = 4.94, p = 0.0263). Similarly to lab males, AB had no impact on starvation resistance in S+P fed wild males (χ^2^ = 1.39, p = 0.2369) (Fig. 5B).

**Figure 5.**
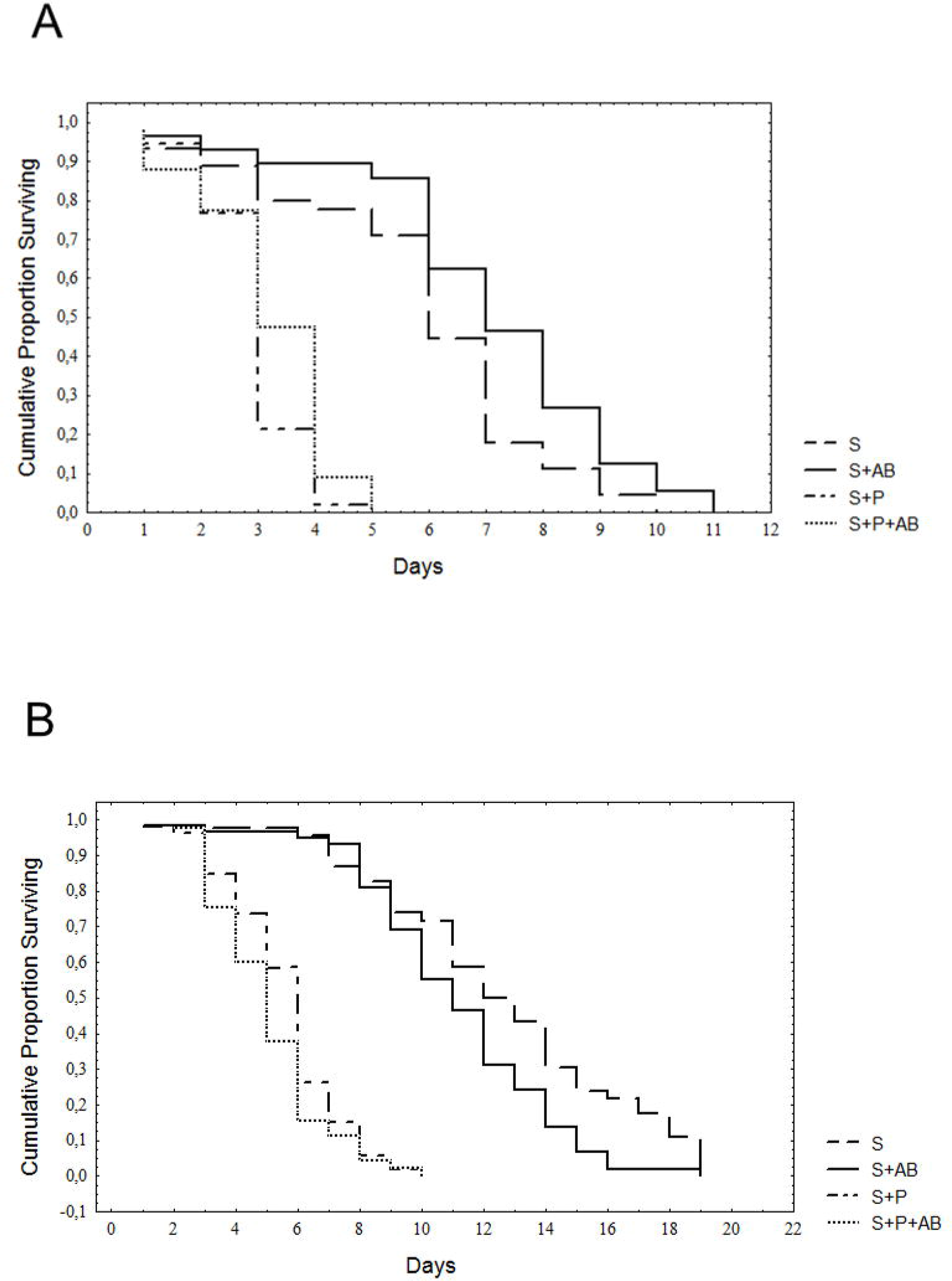
Effect of antibiotics on laboratory (A) and wild (B) *Anastrepha fraterculus* males’ starvation resistance. Cumulative proportion of surviving males fed on S or S+P diets with or without the addition of antibiotics (AB).

### Dry weight

Antibiotics did not affect the adult dry weight both for lab and wild males (F_1,10_ = 1.92, P = 0.1962 for S fed lab males; F_1,10_ = 0.25, P = 0.6263 for S+P fed lab males; F_1,10_ = 0.13, P = 0.7227 for S fed wild males; F_1,10_ = 1.68, P = 0.2235 for S+P fed wild males) (Fig. 6).

**Figure 6.**
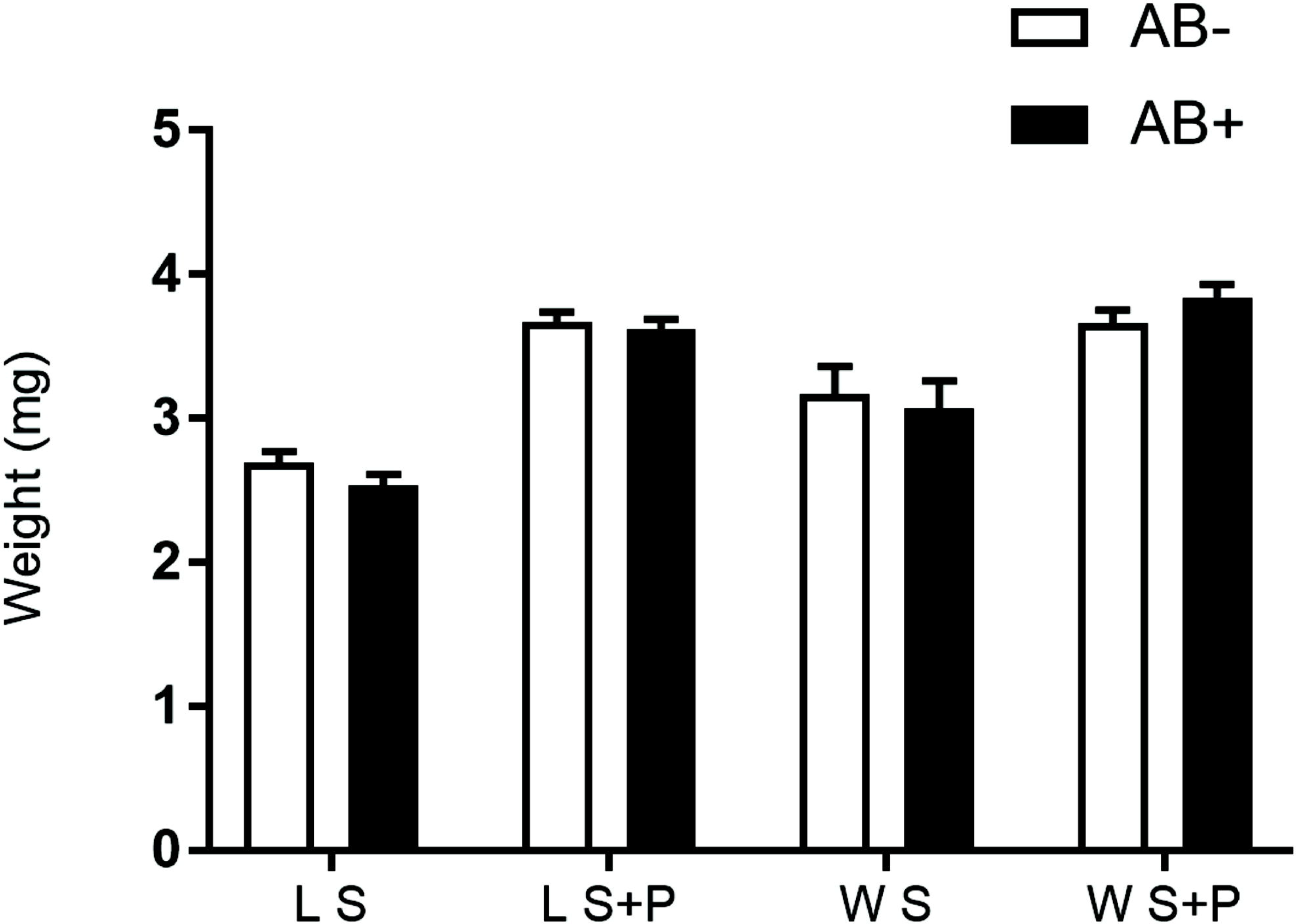
Effect of antibiotics on laboratory and wild *Anastrepha fraterculus* males’ dry weight. Weight (mg) of males fed on S or S+AB and S+P or S+P+AB diets with or without the antibiotic addition (AB).

### Nutritional reserves

Antibiotic treatment had no effect on total sugar content in any combination of male origin and diet (F_1,4_ = 1.19, P = 0.3375 for S fed lab males; F_1,4_ = 3.12, P = 0.1522 for S+P fed lab males; F_1,4_ = 0.001. P = 0.9769 for S fed wild males; F_1,4_ = 1.23, P = 0.3297 for S+P fed wild males) (Fig. 7A). Likewise, AB had no impact on the glycogen content for both origins and type of diets (F_1,4_ = 0.94, P = 0.3876 for S fed lab males; F_1,4_ = 1.35, P = 0.3103 for S+P fed lab males; F_1,4_ = 0.30, P = 0.6144 for S fed wild males; F_1,4_ = 7.23, P = 0.0547 for S+P fed wild males) (Fig. 7B). The analysis of protein content showed a negative effect of AB for S+P fed lab males (F_1,4_ = 53.33, P = 0.002) (Fig. 7C). For the rest of the treatments, no significant differences in protein content were detected between diets containing or not AB (F_1,4_ = 2.90, P = 0.1637 for S fed lab males; F_1,4_ = 0.01, P = 0.9222 for S fed wild males; S+P: F_1,4_ = 0.42, P = 0.5532 for S+P fed wild males) (Fig. 7C). Lipid content was also negatively affected by AB for S+P fed lab males (F_1,4_ = 18.41, P = 0.0127) (Fig. 7D). For the remaining combinations, no differences were found in the lipid content between AB treated and non-treated males (F_1,4_ = 3.62, P = 0.1298 for S fed lab males; F_1,4_ = 0.07, P = 0.8095 for S fed wild males; F_1,4_ = 0.18, P = 0.6938 for S+P fed wild males) (Fig. 7D).

**Figure 7.**
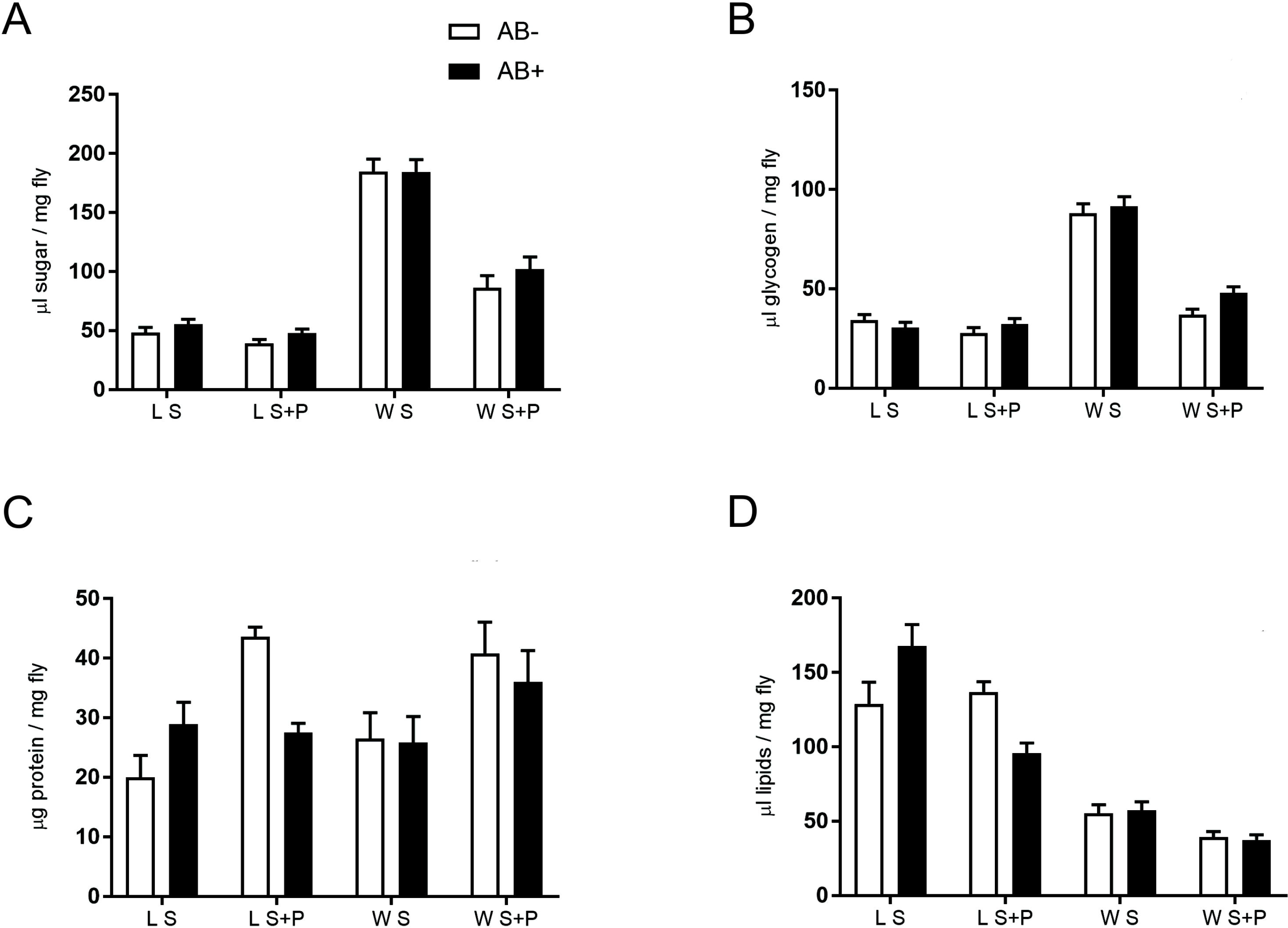
Effect of antibiotic on laboratory and wild *Anastrepha fraterculus* males’ nutritional reserves. (A) Sugar, (B) Glycogen, (C) Protein and (D) Lipids content in males fed on S or S+AB and S+P or S+P+AB diets with or without the antibiotic addition (AB).

## Discussion

Symbiotic bacteria play an important role in the development and biology of many insect species. Recently, an increasing number of studies have focused on the interaction between bacteria and Tephritidae fruit flies [e.g, 5, 14, 19–23, 25, 27]. Our data suggest that bacteria might affect in a positive way several parameters directly related to the mating success of laboratory *A. fraterculus* males, as well as their nutritional status, but would negatively affect their survival under starvation. Specifically, this is supported by the fact that ingestion of antibiotic was associated to detrimental effect in males fed on both types of diet. In S fed males, AB produced a decrease in their sexual display rate, a decrease in the amount of three pheromonal compounds and a mild reduction in mating competitiveness. For S+P males, AB affected the amount of copulas obtained by males, which was correlated with a decrease of protein content. The effect of AB on fitness related parameters depended on two additional factors: the origin of the males (wild or lab) and the presence of a proteinaceous source in the adult diet. Nonetheless, it is important to mention that our results were obtained by an indirect approach under which males received AB as a means of disrupting symbiotic association with bacteria. Even when we found a drastic change in the gut microbiota, and we associated this with a reduction of the overall fitness of the males, AB could have also affected the mitochondria [50] causing (or at least contributing to) a decrease in mating success and related parameters. This is a limitation of the current experimental approach and should be considered in further studies, for example by inoculating specific bacteria to the diet. This approach has shown promising results in different fruit fly species, such as *Dacus ciliatus* (Loew) [51], *C. capitata* [10, 27, 30, 31] and *B. oleae* [29].

### Analysis of the gut’s bacterial community and the effect of antibiotic treatment

We found that the incorporation of AB in the adult diet affected the bacterial community of the digestive tract of *A. fraterculus* males. Similar results were obtained for another fruit flies like *C. capitata* and *B. oleae* subjected to similar antibiotic trials [5, 19–23]. In our experiments, the presence of AB had no impact on the decision to feed on a given food source. This shows neither a phagostimulant nor a deterrent effect of adding AB into the diet. DGGE followed by sequencing showed a dominant representation of the Enterobacteriaceae family in the *A. fraterculus* male gut, as has been previously evidenced for other fruit fly species (see [52] for a review). Some of these microbial taxonomic groups are composed by diazotrophic bacteria (i.e., nitrogen fixers) with an essential function in the acquisition of nitrogen compounds and carbon metabolism, allowing both sexes to reach their reproductive potential [12, 13, 53–55]. The strong impact of AB on potentially key symbiotic bacteria evidenced in males, suggest a similar approach could provide relevant information on the role of gut bacteria in females as well. Antibiotics appear to have drastically affected the gut enterobacterial diversity, since other taxonomic classes (e.g., *Klebsiella sp., Enterobacter sp*. and *Serratia sp*.) were not detected in adult males’ flies under AB treatment. These differences in the gut bacterial community found between AB-treated and non-treated individuals were also supported by the linkage dendrogram analysis of DGGE profiles. This reduction in gut bacterial diversity, associated to physiological changes in the host has been previously reported for Tephritidae fruit flies [5, 19, 20, 21] as well as for other insect species [56].

### Impact of antibiotic treatment on reproductive parameters, nutritional status and starvation resistance

*Anastrepha fraterculus*, similarly to other tephritid species, presents a lek based mating system [43, 57] in which males aggregate and perform sexual displays (calling behavior) to attract females to a mating arena that has neither resources nor refuges [58]. The sexual display involves acoustic, chemical and visual signals (e.g., wing fanning, the extrusion of the salivary glands and protrusion of the anal tissue) [59], and is therefore an energetically demanding task ([60], reviewed in [61]). This means that adults need to acquire specific nutrients in order to complete their sexual development [54, 61, 62]. Numerous studies have found that protein intake has a positive impact on the reproductive success of *C. capitata* males, affecting their ability to participate in leks [63], to emit pheromone [64, 65], to transfer a substantial ejaculate [66] and to decrease female receptivity [67]. In the same way, studies with other *Anastrepha* species showed that protein intake results in an improvement of male’s sexual competitiveness [46, 62, 68, 69], as well as an increase in the amount of pheromone released by males [70]. In the present study we found significant differences in the amount of lipids and proteins between lab males that were fed with AB and those that were not, for S+P treatment. For both nutrients, the addition of AB to the diet had a negative effect on the nutritional reserves compared to males that retained their gut bacteria. The effect of AB on the nutritional reserves of S+P fed lab males correlates with a significant decrease of the amount of copulas reached by these males compared to non-treated males. Ben-Yosef et al. [19] also observed for S+P fed males a decrease (although not significant) in the reserves of protein after the addition of AB and an impact on mating related variables (see below).

The higher mating competitiveness in S+P fed non-treated lab males was not associated to higher rates of sexual displays or sex pheromone emission. Henceforth, it seems that females were able to assess the nutritional status of the males, in spite of the lack of differences in these components of the courtship, maybe using more subtle, close range signals that were not recorded in this study. For several tephritid species, acoustic communication has major implications on mating success. For instance, in several *Anastrepha* species the sound produced by repeated bursts of wing-fanning generates pulse trains that stimulate the females [71–75]. Likewise, behavioral male-male or male-female interactions (e.g., movements, fights or contacts) could be influencing female choice [59]. In our case, females could have used multiple signals to assess males’ quality, rejecting those of poor quality related to a low amount of protein as result of a change in their gut bacteria community [52]. Alternatively, males with larger reserves could be more aggressive in defending small territories, a parameter that was not assessed in our experiments. Observations at a finer scale (like video or sound recordings) and also at a higher scale (like field cages with host trees inside) may help to reveal the targets of female choice that could be affected (directly or indirectly) by gut bacteria.

Several studies tested the hypothesis that bacteria contribute to mating success of *C. capitata*. Most of them followed a direct approach adding specific bacterial strains as probiotics into artificial diets and showed an increase in male mating success [27, 30, 39] with some exceptions [25, 31]. Ben-Ami et al. [39] found that irradiation of *C. capitata* pupae affected the abundance of adult gut bacteria, more specifically *Klebsiella oxytoca*, and this was associated to a reduction of male mating success. Following an indirect approach, as the one used in the present study, Ben Yosef et al. [19] found that *C. capitata* males that were fed antibiotics needed more time to mate (higher latency times) than males that did not received antibiotics, and only when the diet contained protein, as no effect of antibiotics was detected for sugar fed males. According to the same study, bacteria could be involved in the production of a more attractive sexual signal (not analyzed), which may have been mediated by a protein-bacterial interaction [19]. This study on *C. capitata*, and the results of the present one on *A. fraterculus*, showed that the manipulation of symbiotic bacteria in S+P fed males affected their nutritional reserves, and this was associated with a decrease of their mating competitiveness, although the precise mechanism by which females respond to these changes is still unknown and differences in the variable in which this was expressed (i.e., latency or mating percentage) can be attributed to differences in the species under study.

Antibiotic treatment also affected parameters associated to the sexual behavior of S fed *A. fraterculus* lab males. For these nutritionally stressed males, AB significantly decrease the rate of sexual displays (wing fanning and exposure of salivary glands) and the amount of three antennally active compounds of the male sex pheromone. Additionally, AB treated males fed on sugar obtained numerically less copulas than non-treated males, even though the differences were not statistically significantly. However, in this case there was no significant difference in any of the analyzed nutrients. Although bacteria do not seem to impact on the nutritional status of S fed males when lipids, carbohydrates and protein were measured, they still could be contributing with other essential nutrients that allow fruit flies to fill ‘deficiency gaps’ (*sensu* [52]) or even to certain essential aminoacids. For example, Ben-Yosef et al. [5, 21] found that the fecundity of females was significantly enhanced by the presence of gut bacteria when flies were fed with a diet containing only non-essential amino acids. This hypothesis needs further research, as it may help to better understand the role of bacteria and even try to supplement artificial diets with specific nutrients as to improve flies’ quality with pest management purposes. In any case, through an indirect approach (i.e., antibiotic treatment) it was possible to observe the benefits of symbiotic bacteria in males fed on poor diets.

When nutritional reserves and parameters associated to the sexual success of *A. fraterculus* were analyzed in wild males, no significant differences were found. However, the addition of AB resulted in a lower, but not statistically different, protein content in S+P fed males, which is similar to what was observed in lab males. It was also observed that the total amount of sugar and glycogen in wild males was much higher in comparison to lab males, which showed larger lipid reserves. All these results showed that removal of gut bacteria (mainly Enterobacteria) at the adult stage was not strongly connected to changes in the nutritional status or mating competitiveness in wild males. This could be the result from at least three different reasons. First, wild males and bacteria could establish an association more similar to a commensalism than to a mutualistic one, being bacteria the only organisms obtaining a benefit, at least when mating is considered. Second, wild flies used in this study had developed in guavas (a primary host for *A. fraterculus*) where the pupal weight is higher than in alternative hosts, such as peach or plum [46]. Guava fruit could provide exceptional nutrients that allow males to reduce the impact of unfavorable conditions, such as the removal of the intestinal microflora. Third, wild flies were provided with an artificial adult diet, which could represent a huge shift compared to natural food sources. This change in environmental and nutritional conditions, associated to the adaptation of wild individuals to artificial rearing conditions, could have produced instability in the microflora constitution and/or a physiological impact on males, adding further complexity and even diluting the contribution of bacteria.

Regarding males’ ability to endure starvation, we found that the effect of AB depended on the type of diet as well as the origin of the males. First, the starvation resistance of S fed males was higher (i.e., lived longer) than S+P fed males, regardless of the addition of AB and the origin of the flies. Similar results were also observed in previous works [61, 64, 68, 76] where adding protein in the diet (although it increased the sexual performance of males), negatively affected their ability to endure starvation [61]. Second, AB had contrasting results for wild and lab males. While for S fed wild males the presence of bacteria gave males a significant advantage over males fed with AB, the addition of AB allows S fed lab males to significantly live longer than males that were not treated with AB. Ben-Yosef et al. [20] also showed that AB treatment positively affects the longevity of males and females fed on sugar. As mentioned before, nutritionally stressed lab males without their gut bacteria (i.e., S+AB males) were found to perform significantly less sexual signaling than S males (and therefore did not spend great amounts of energy), which may have leave them in better nutritional conditions to endure starvation. Alternatively, the addition of AB could have removed pathogenic bacteria which could be more widespread in laboratory due to the rearing conditions [39]. For example, Behar et al. [22] found that inoculation of sugar diet with *Pseudomonas aeruginosa* reduced the longevity in *C. capitata*.

## Conclusions

In summary, following an indirect approach (AB treatment) potential contributions of the gut bacteria associated to *A. fraterculus* males was found. These contributions to the fitness of the male were more evident for laboratory flies fed on sugar and protein. This could be mediated by a combination of higher protein reserves and bacteria presence in S+P diets, which leads to a greater male competitiveness; whereas the absence of protein and presence of bacteria in S diets does not improve nutritional reserves but increases the rate of sexual displays, the amount of pheromone emitted and enhances the sexual success of the males. Thus, the evidence suggests that gut microbiota includes beneficial bacterial species that are able to exert a positive contribution. Removal of bacteria had nonetheless a positive effect on starvation resistance in sugar fed lab males, which probably points out to the presence of pathogenic strains in the rearing or the inability of sugar fed to cope with the energetic demand associated to reproduction, or both. Our results have important implications for the development and effectiveness of SIT for *A. fraterculus* although the role of gut bacteria should be confirmed following a more direct approach (i.e., the addition of specific bacterial strains to the diet). Likewise, the characterization of the gut bacterial community associated to females and its potential impact throughout the life cycle should be further addressed.

## Materials and methods

### Biological material and holding conditions

Experiments were carried out with wild and laboratory-reared *A. fraterculus* flies of the Brazilian-1 morphotype. Wild pupae were recovered from infested guavas (*Psidium guajava* L.) collected at Horco Molle, Tucumán, Argentina. Laboratory flies were obtained from the colony held at INTA Castelar. Rearing followed standard procedures [77, 78] using an artificial diet based on yeast, wheat germ, sugar, and agar for larvae [79] and a mixture of sugar and hydrolyzed yeast (MP-Biomedical®, Santa Ana, California, USA) (3:1 ratio) for adults. Rearing was carried out under controlled environmental conditions (T: 25 ± 2°C, RH: 70 ± 10%, photoperiod 14L: 10D) until adult emergence.

### Diets and antibiotics

Males from the two origins (wild or lab) were provided with one of two different diets: sugar (S) or sugar + hydrolyzed yeast (S+P), which in turn could have been supplemented or not with antibiotics (AB). This procedure resulted in four treatments: 1) S; 2) S+AB; 3) S+P; 4) S+P+AB. The S+P diet consisted of 3:1 mixture of sugar and hydrolyzed yeast, which constitutes a rich source of peptides, amino acids, vitamins and minerals, in addition to carbohydrates [5] and is comparable with artificial diets that provide the flies with all their nutritional needs [19, 20, 80]. Because we aimed at comparing the impact of AB between males that had access to protein sources and males that were deprived of protein, S diet was supplemented with NaCl, MgSO_4_, H_3_BO_3_ and a complex of vitamins (A, D, B1, B2, B3, B5, B6, B9, B12, C) and minerals (FeSO_4_, Ca_3_(PO_4_)_2_, CuSO_4_, Ca(IO_3_)2.6H_2_O, CoSO_4_, MnSO_4_, MgSO_4_.7H_2_O, ZnSO_4_, Mo, K_2_SO_4_) (DAYAMINERAL, Laboratorios Abbot, Buenos Aires, Argentina). This way, S and S+P diets were as similar as possible in terms of micronutrient content. AB treatment consisted of Ciprofloxacin (10µg mL^-1^) and Piperaciline (200 µg mL ^−1^), which proved to be the most potent antibiotic combination for the inhibition of bacterial growth in *C. capitata* [19]. The different components of each diet were mixed with distilled water to form a liquid diet. For most experiments, the diet solution was applied to a piece of filter paper and placed inside the cages, and replaced every 48 h. Only when consumption was evaluated (see below), the diets were placed in a container (the lid of a 2 ml Eppendorf vial) and left inside the cage. The diets were colored with a food dye (FLEIBOR, Laboratorios Fleibor, Buenos Aires, Argentina) to allow the differentiation between those males that had been fed with AB and those that had not. This marking system does not present any detrimental effect on *A. fraterculus* [48, 81].

### Intake of antibiotic supplemented diets and its effect on gut bacteria diversity

#### Diet consumption

To evaluate whether the presence of antibiotic affected the rate of food consumption, males were offered either S and S+AB diets, or S+P and S+P+AB diets in a dual choice experiment. For each male origin and type of diet, three replicates were evaluated. In each replicate, 20 recently emerged males (< 24-h old) were confined in a 1 L plastic container and provided with diets as a solution (500 µl of initial volume – V_0_) placed in two different vials. Diet consumption was determined every 48 h by removing the vials containing diet and measuring the remaining volume of diet (Vr) with a Hamilton syringe. For each recording, the volume consumed (Vc) was calculated as: V_0_ - Vr + Ve (the volume of diet lost due to evaporation). Ve was estimated from control vials which contained the different diets but no flies. Every time a vial was removed for measuring Vr, a new vial with 500 µl of diet was placed in the cage. The number of flies that remained alive at each recording was used to estimate individual consumption (Vci) during the 48 h time interval in which the vial was exposed (Vci = Vc/number of individuals alive in the cage). The experiment lasted 18 days, and the Vci from subsequent 48 h periods were added to obtain the total individual consumption (Vti).

#### Molecular characterization of gut bacteria

Ten-day-old virgin males from each origin, type of food and treatment were washed 3 times in ethanol 70% and their guts were dissected. Total DNA from single fly guts was extracted following Baruffi et al. [82] protocol with some modifications of volume due to the size of the tissue under study (gut of individual fly), and used as template to amplify the V6-V9 variable region of the bacterial 16S *rRNA* gene by PCR and posterior DGGE fingerprinting, using the primers 968F-GCclamp / 1408R [83].

DGGE was conducted using a DcodeTM system (Bio-Rad) and performed in 6% polyacrylamide gels, containing 37.5:1 acrylamide:bisacrylamide and a denaturing gradient of 35:70% and 40:60% of urea. The gels were stained for 30 min in 1X TAE buffer containing ethidium bromide and visualized in a UV trans-illuminator. DGGE marker was prepared from a selection of bacterial 16S *rRNA* gene products to enable gel to gel comparison. For the identification and subsequent characterization of DGGE bands, a selection of bands was made according to their position in the electrophoretic profiles. This selection included bands that were shared between individuals (located at the same position in different lanes) and some others that were exclusively present in one individual (differentially located), in order to get a representative sampling of all bands in the DGGE profile. DGGE fragments of interest were numbered and excised with sterile razor blades immediately after staining and visualization of the gels. Gel bands were stored in 50 µl distilled water at –20ºC and eluted at 4°C overnight before PCR reaction. DNA was reamplified using the PCR-DGGE primers without the clamp, and product integrity was checked by agarose gel electrophoresis. The PCR products were purified using the QIAGEN PCR purification kit (Qiagen Ltd, Hilden, Germany) and directly sequenced with 968F primer.

V6-V9 (approximately 440 bases) 16S *rRNA* gene sequences obtained from DGGE bands were aligned using BioEdit [84] and Clustalw [85]. Sequence similarity searches were performed using the online sequence analysis resources BLASTN [86] of the NCBI (nt database) and Seqmatch provided by the Ribosomal Database Project (RDP) [87]. Alignment of our sequences and the closest related taxa was carried out using the MEGA 6.06 software package. A phylogenetic tree based on distance matrix method was constructed. Evolutionary distances were calculated using the method of Jukes and Cantor [88] and topology was inferred using the ‘‘neighbor-joining’’ method based on bootstrap analysis of 1,000 trees. Phylogenetic tree calculated by maximum parsimony using the PAUP phylogenetic package was also generated.

Nucleotide sequences generated from 16S *rRNA* gene corresponding to *A. fraterculus* gut bacteria, and obtained from DDGE purified bands, were submitted to GenBank (https://www.ncbi.nlm.nih.gov/genbank/index.html‎). The samples were named as follows: 1 S+P+AB Wild; 10 S+P+AB Wild; 4 S+P Wild; 5 S+P Wild; 6 S+P Wild; 5 S+AB Wild; 3 S Wild; 1 S+P+AB Lab; 2 S+P+AB Lab; 5 S+P Lab; 3 S+P Lab; 4 S+P Lab; 4 S+AB Lab and 5 S Lab. The corresponding accession numbers are: MH250014-27, respectively.

### Impact of antibiotics on reproductive parameters

#### Males’ mating competitiveness

To evaluate males’ mating competitiveness, one wild sexually-mature virgin female (14 days-old) was released inside a mating arena (a 1 L plastic container), which contained two males from the same origin as well as diet, but only one had received AB. Males were fed on the diets from emergence until sexual maturity (14 days-old), time at which they were tested. After the female was released in the arena, the occurrence of mating was followed by an observer. The type of male, the copula start time and the time at which flies disengaged were recorded. The experiment was conducted under laboratory conditions (T: 25 ± 1ºC and 70 ± 10 % RH) from 8:00 to 11:00 am. The experiment was replicated on different days as follows: five days for wild males (both S and S+P diets), three days for S fed lab males and four days for S+P fed lab males. We evaluated 667 mating arenas: 191 for S fed wild males and 171 for S+P fed wild males, 145 for S fed lab males, 160 for S+P fed lab males.

#### Males calling behavior and chemical profile

To evaluate the potential changes in male sexual signaling related to the AB treatment, males’ calling behavior was recorded at the same time that male-borne volatiles were collected. Each replicate consisted of ten males from the same combination of diet and AB treatment, placed in a 250 mL glass chamber (20 cm length, 4 cm in diameter) [81]. Males were 10 days-old and were kept under the aforementioned treatments until the day of the test. Eight replicates were carried out and only lab males were analyzed.

Behavioral recordings and collection of volatiles started at 8:30 am and lasted for 3 h [daily period of sexual activity for this *A. fraterculus* morphotype (43)]. Two components of male courtship associated with pheromone emission and dispersion were considered: wing fanning and exposure of salivary glands [43, 59, 89]. During the observation period, the number of males performing these behaviors was recorded every 30 minutes. At the same time, the volatiles emitted by the calling males were collected by blowing a purified air stream through the glass chambers. Volatiles were collected onto traps made of 30 mg of Hayesep Q adsorbant (Grace, Deerfield, IL, USA) [81]. After collection, the trapped volatile compounds were eluted with 200 µl of methylene chloride and chemically analyzed using an Agilent 7890A gas chromatograph (GC) equipped with a HP-5 column (30 m ± 0.32 mm inner diameter ± 0.25 µm film thickness; Agilent Technologies), and an Agilent 5977 mass spectrometer. The initial oven temperature was 35°C and after 1 min the oven temperature was increased to 100°C at 5 °C min^-1^ and from 100°C to 230°C at 12°C min^-1^, then held for 10 min. Samples were injected in the splitless mode with the injector purged at 30 s with helium as the carrier gas at 27.6 cm/sec flow velocity. Methyl nonadecanoate (5 ng per 1 µl of methylene chloride) was used as internal standard. The compounds were identified by using their relative retention times and comparison of their mass spectra with libraries. The identity of specific compounds (e.g., limonene, suspensolide, (E,E)-α-farnesene, anastrephin and epianastrephin) was also confirmed with standards.

In order to analyze the effect of AB on the chemical profile of the cuticle, after the pheromone sampling ended males were gently removed from the glass chambers and washed (in groups of ten) with 1 ml of hexane for 1 min in 2 ml glass vials. Methyl nonadecanoate (5 ng per 1 µl of hexane) was used as internal standard. Compounds were identified as described above.

### Impact of antibiotics on starvation resistance and nutritional status

#### Starvation resistance

To evaluate the effect of antibiotics on males’ ability to endure starvation, a group of 20 wild or lab males (< 24-h old) was caged in a 1 L plastic container and fed one of the aforementioned diets. Food was replaced every 48 h. After 10 days, food was removed and only water was provided. Every 24 h, the number of dead males was recorded until all individuals had died. For each origin and treatment, the experiment was replicated three times.

#### Dry weight and nutritional reserves

To evaluate the effect of AB on males’ dry weight and nutritional reserves, groups of 20 wild or lab males (< 24-h old) were placed in 1 L plastic containers and provided with one of the aforementioned diets (i.e., S; S+AB; S+P; S+P+AB). Six cages were arranged per diet and origin. Diet was replaced every 48 h. After 14 days, males were removed from the cage and preserved at –20°C. A sample of 10 individuals from each cage was dried out in an oven at 50°C for 5 h and weighed in a precision scale (readability: 0.0001 g) (Ohaus Corporation, Parsippany, NJ, USA). Nutritional reserves were determined following standard biochemical techniques. Protein content was determined with the Bradford [90] method using Coomassie brillant blue G-250 reagent. Lipid and carbohydrate contents were determined with the Van Handel [91] method. Total sugar and glycogen contents were measured with anthrone reagent [92] whereas vanillin reagent was used for lipid measurement [93].

### Statistical Analysis

Data were analyzed using InfoStat and R software [94, 95].

To determine whether the presence of AB in the diet affected diet consumption, a mixed effect model analysis for each combination of diet and origin was performed with AB treatment as the fixed factor and the cage from which the flies were taken as the random factor.

To evaluate the AB effect on the percentage of copula achieved by treated and non-treated males, we performed a mixed effect model analysis with AB treatment as the fixed factor and the day of the experiment as the random factor. After verifying lack of heteroscedasticity, the data were analyzed without transformation. For wild males fed on the S diet, two experimental days (replicates) were removed due to the low number of copulations recorded (less than 10 matings). Latency was analyzed by Mann-Whitney test for each category (male origin and diet regimen) separately. Copula duration was analyzed with a mixed effect model where the fixed factor was the AB treatment and the random factor was the day of the experiment.

The mean number of males exposing their salivary gland or fanning their wings across the observation period was compared between S and S+AB, or S+P and S+P+AB, by means of Student’s *t*-tests. The abundances of volatile and cuticle compounds were obtained by computing the ratio between the area under the peak of each compound and the area under the peak of the internal standard. Then, the abundance of each compound was compared between AB treated and non-treated males (separately for S and S+P males) in two ways. First, a Student’s *t*-test was performed for each single compound detected by the mass detector. Second, a new Student’s *t-*test was performed by building a new variable that resulted from adding those compounds that showed evidence of electroantennal activity in *A. fraterculus* females of the same laboratory strain we used in this study. These compounds included: E-β-ocimene; Z-E-α-farnesene; E-E-α-farnesene; and epianastrephin [96, 97].

To evaluate the effect of AB on starvation resistance, the data were analyzed using a Kaplan-Meier survival analysis for each male origin and diet combination separately. The effect of AB on males’ dry weight and nutritional reserves were analyzed by means of mixed effects models in which the AB treatment was the fixed factor and the cage from which the flies were taken was the random factor.

## Acknowledgments

Authors would like to thank Fabian Milla and Germán Crippa for supplying laboratory flies. Authors are grateful to the editor and two anonymous reviewers for their constructive comments, which helped us to improve the manuscript.

## Abbreviations

AB: Antibiotic; SIT: Sterile insect technique; S: Sugar; S+P: Sugar + hydrolyzed yeast; DGGE: Denaturing gradient gel electrophoresis; RDP: Ribosomal database project; UPGMA: Unweighted pair-group method with arithmetic averages; GC: Gas chromatograph.

## Ethics approval

Not applicable.

## Consent for publication

Not applicable.

## Availability of data and materials

All data generated or analyzed during this study are included in this published article (and its supplementary information files).

## Author contributions

MLJ, MJR, LG and MTV performed the diet consumption, mating competitiveness, dry weight, nutritional reserves and starvation resistance tests. LEP and SBL conducted the molecular analysis. DFS, GEB and PCF carried out the calling behavior, and volatile and cuticle compounds analyses. PMP and FC performed the nutritional reserves analyses. DFS, SBL, KB, JLC and MTV conceived the project and coordinated the activities. MLJ, LEP, SBL, PCF, JLC, KB, MTV and DFS wrote the paper. All authors interpreted the results and commented on the manuscript. All authors read and approved the final manuscript

## Additional files

Additional files 1:Figure **S1**. Alignment of V6-V9 16S *rRNA* nucleotide sequences (420 bases) obtained from DGGE profiles. Af V6-V9 Seq 1-14 correspond: Band 1 S+P+AB Wild, Band 10 S+P+AB Wild, Band 4 S+P Wild, Band 5 S+P Wild, Band 6 S+P Wild, Band 5 S+AB Wild, Band 3 S Wild, Band 1 S+P+AB Lab, Band 2 S+P+AB Lab, Band 5 S+P Lab, Band 3 S+P Lab, Band 4 S+P Lab, Band 4 S+AB Lab, Band 5 S Lab respectively (WORD 51.5 KB).

